# Stem embolism vulnerability curve depends on methods used: is there a fifth mechanism of cavitation?

**DOI:** 10.1101/2022.01.05.475122

**Authors:** Guoquan Peng, Lei Cao, Zhiyang Ren, Zhao Liang, Guo Yu, Dongmei Yang, Melvin T. Tyree

**Affiliations:** College of Chemistry and Life Sciences, Zhejiang Normal University, Jinhua 321004, China

**Author notes:** **Corresponding author:** Dongmei Yang, College of Chemistry and Life Sciences, Zhejiang Normal University, Jinhua 321004, China. **Author contributions**, D.Y. and M.T. conceived the research plans; G.P. and D.Y. supervised the experiments; G.P. performed most of the experiments and Z.L. performed staining experiments on *Robina* in Yangling; L.C., Z.R and G.Y. performed the experiments for species in Jinhua. D.Y. and G.P. analyzed data and wrote the manuscript; M.T. revised the manuscript. D.Y. agrees to serve as the author responsible for contact and ensures communication. **Responsibilities of the author for contact**, The author responsible for distribution of materials integral to the findings presented in this article in accordance with the policy described in the Instructions for Authors (https://academic.oup.com/plphys/pages/General-Instructions) is Dongmei Yang.

**Keywords:** vulnerability curves, nano-particles, *Robinia pseudoacacia*, centrifuge techniques, recalcitrant curves, origin of nano-particles

## Abstract

A long-established ecological paradigm predicts a functional relationship determining vulnerability to cavitation: vulnerability increases with vessel hydraulic efficiency and vessel diameter. Even within a species, big vessels cavitate before small ones.

Some centrifuge methods for measuring vulnerability are prone to artifacts due to nano-particles seeding early embolism, as the particles are drawn into vessels during measurements. Both the Sperry and Cochard rotors are prone to early cavitation due to nano-particles drawn into long and wide vessels in *Robinia pseudoacacia* and *Quercus acutissima*, whereas extraction centrifuge methods produce vulnerability curves more resistant to cavitation.

Sufficient nano-particles pass through the stems to seed early embolism in all rotor designs. For several years, people have thought that early embolism is induced by nano-particles present in laboratory water. One new hypothesis is that the origin of nano-particles is from cut-open living cells but a much bigger study including many species is required to confirm this idea. This paper confirms the hypothesis in comparisons between short-vesselled *Acer*, and long-vesselled *Robinia, and Quercus*. Our new results and a review of old results justifies bigger study.

Hypothetical nano-particles might explain why different methods for measuring vulnerability curves cause different *T_50_* = tensions causing 50% loss of hydraulic conductivity. Hence the hypothesis for future research should be that the open-vessel artifact is consistent with ‘long’ vessels surrounded by cut open living cells.

**One sentence Summary:** Nano-particles induced early cavitation in species with vessel lengths about ¼ the stem length used in all centrifuge rotors, and the origin of nano-particles might be from living cells nearby vessels

## Introduction

Water transport in land plants is fundamentally unstable, and the results in this paper and the recent literature enhance our understanding of the structural paradigms driving the evolution of xylem anatomy that permits metastable water transport. The Cohesion-Tension Theory advanced by Dixon and Jolly (1897) proposed that water is transported in a quasi-stable tensile status (Tyree, 1997). The concept of liquids having a tensile property is anathema to mechanical engineers and physicists because solids, by definition, have tensile properties, but liquids, by definition, have no tensile properties; nevertheless, plants seem to succeed in tensile water transport down to negative pressures of −12 MPa, but more typically in the range of −1 to −3 MPa (Tyree and Zimmermann, 2002; Cochard et al., 2013). Loss of hydraulic conductivity, *K_h_*, in stems occurs because of cavitation-induced water loss from xylem conduits. The tensile strength of water in xylem is measured in terms of the tension required to reduce hydraulic conductivity by 50%, *T_50_*. In this paper *T* is positive having pressure units equal to *-P*, where *P* is the absolute pressure of water. Hence water at 0.1 MPa absolute pressure (water in a beaker) has a tension of −0.1 MPa, water held in a perfect vacuum has a tension of 0 MPa and water at −1 MPa absolute *P* has a *T* = +1 MPa

It has long been assumed that centrifuges provide the fastest and most reliable way of measuring *T_50_*. Fully hydrated stem segments are placed inside a specially designed rotor and spun in a centrifuge to induce embolism where the maximum tension, *T_max_*, occurs at the axis of rotation and falls in a bell-shaped curve (Fig. 1) to zero at two water surfaces where water held by cuvettes. The stems are supported by rotors with their ends emersed in water. However persistent questions have arisen concerning what is called the long-vessel artifact that seems to result in *T_50_* that are arguably too low, compared to *T_50_* measured by other techniques (Cai et al., 2014; Wang et al., 2014; Yin et al., 2018; Du et al., 2019; Peng et al., 2019). There are two kinds of rotors in use for spinning in different models of centrifuges: (1) The traditional Sperry rotor is mounted in a floor-model centrifuge and can spin 3 stem segments simultaneously to induce embolism, but embolism, measured through loss of *K_h_*, has to be measured outside the centrifuge (Alder et al., 1997), which is sometimes called the static centrifuge method; and (2) the Cochard rotor that can spin only one stem segment at a time in a smaller table-top centrifuge but measurements of *K_h_* can be performed while the stems spin (Cochard et al., 2005), hence, it is an *in situ* flow technique. These centrifuges are capable of rapidly and precisely measuring vulnerability curves, VC, which are plots of % loss of conductivity versus *T* at the axis of rotation. The long vessel artifact occurs in species with mean vessel length approaching the half-length of the stems being spun in a rotor (rotor diameter approximately 0.14 to 0.28 m long). The artifact produces vulnerability curves, VC, which have *T_50_* at lower values than that measured by slower but more traditional methods, such as desktop dehydration of large shoots (1 to 2.5 m long) and by the staining methods.

**Fig. 1.**
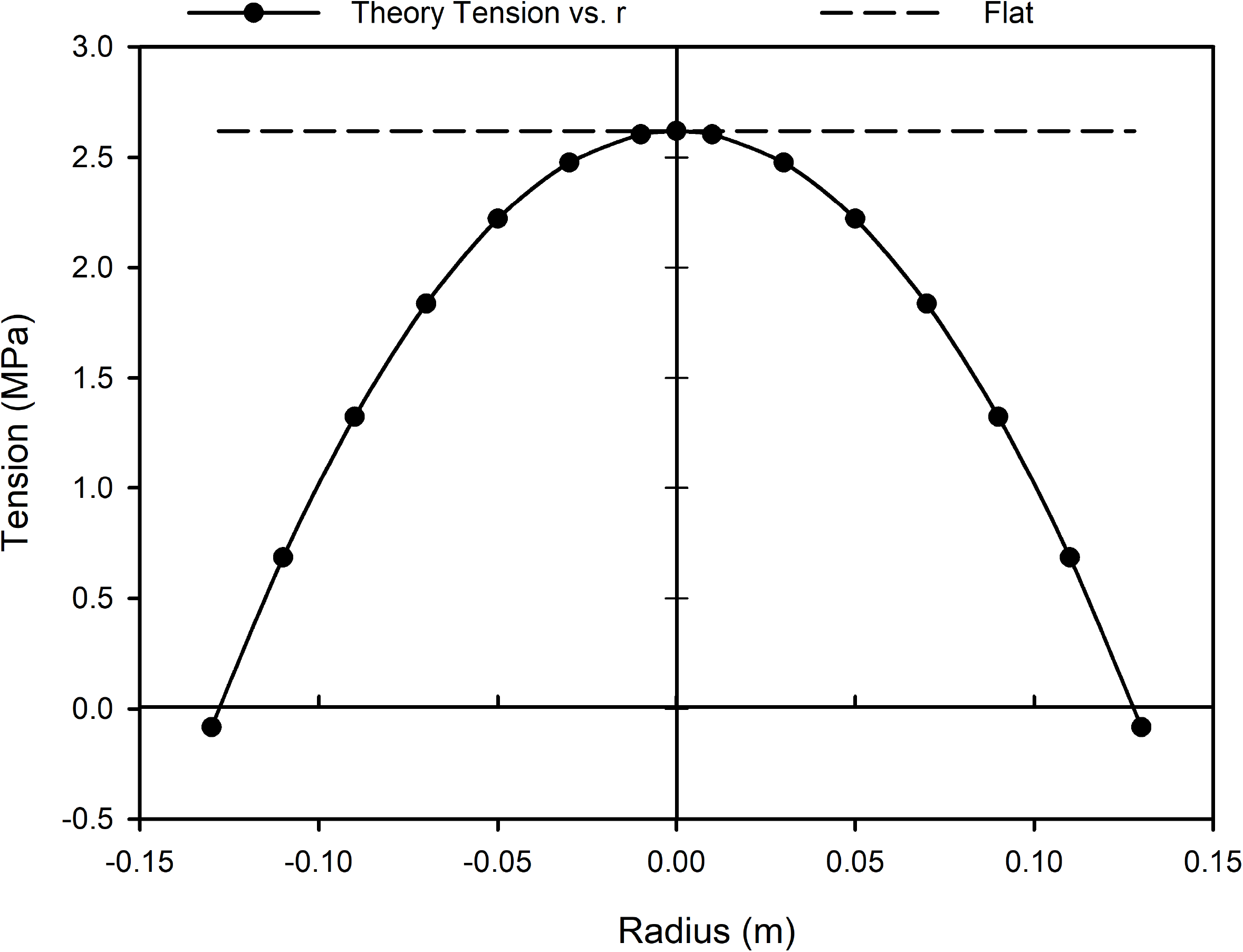
The theoretical profiles of tension in stems versus stem position. Solid line with circles = tension versus position relative to the center of spin of a centrifuge rotor. The flat dashed line is the tension versus similar distance during benchtop dehydration of an excised shoot.

The only way the water can sustain tension is when it is enclosed in some kind of pipe or chamber without an air space, i.e., only chamber walls and water. But the nature of the “chamber”, for example, a glass tube versus a plant vessel, makes a difference to the stability of the system before it cavitates and the water breaks to form two distinct liquid and gas phases. So far only four mechanisms of cavitation have been proposed: (1) Homogeneous nucleation inside a body of water (Briggs, 1950; Pickard, 1981), which occurs at about *T* ≈ 100 MPa. (2) Heterogenous nucleation at the surface of water filled-glass tubes curved into an s-shape and spun in a centrifuge (Briggs, 1950). In glass, cavitations occur at about *T* ≅ 25 MPa. (3) Adhesive failure at the surface cellulose vessel walls (Pickard, 1981; Tyree, 1997). (4) Air seeding at pit membranes between an embolized vessel adjacent to a water-filled vessel which occurs at −1 to −12 MPa and is often assumed to be the dominant mechanism in plants. In pits the air seeding is viewed as occurring in the irregular porous spaces between cellulose fibers of the pit membranes. Pit membranes occur 100 times per m in a short vessel (0.01 m long) and less frequently in long vessels >0.2 m up to 1 or more m long (Tyree and Zimmermann, 2002). The “tensile” strength of these porous spaces is thought to be related to the surface tension of water, *τ*, which can support an air-water interface with a radius of curve given by *T* ≈ 2*τ/r*, where *r* is the effective radius of curvature of water. A value of *T* = 1 to 10 MPa can be supported by a radius of curvature of 100 to 10 nm, respectively.

This paper addresses the recent literature about ‘long vessel artifacts’ wherein the *T_50_* measured on stems of long-vessel species in a centrifuge is substantially smaller then *T_50_* measured outside the centrifuge, e.g., bench top dehydration of shoots > 2 times longer the maximum vessel length. The study is performed in more precise centrifuges with advanced temperature control available in Chinese-built centrifuges to obtain more consistent results

The value of *T_50_* tends to decline with increasing vessel diameter or vessel surface area (Hacke et al., 2006), and vessel length tends to increase with vessel diameter (Cai and Tyree, 2010; 2014). Until recently, the dominant hypothesis to explain these relationships is the air-seeding hypothesis of the mechanism of embolism via pit pores. The air seeding hypothesis implies that once some vessels have embolized at a given tension, *T_i_*, that no more embolisms will occur until a higher tension, *T_f_* > *T_i_* is applied. It is already known that repeated measurement of *K_h_* in a Cochard rotor at the same tension induces more loss of *K_h_* (Wang et al. 2014).

In this paper we will discuss previous literature on the open-vessel artifact and then examine the merits of a 5^th^ hypothesis for embolism. We propose that nano-particles seed at least some cavitations at tensions below the tension causing embolisms via air seeding at pit membranes. But this happens only in cut open vessels, because the proposed nano-particles are too large to pass from soils to fine roots in intact plants and too big to pass through pit membranes between adjacent vessels in cut branches. We also propose that new nano-particles are introduced after each cycle of spinning in a Sperry-type rotor then measuring *K_h_* in a low-pressure flowmeter, or when injecting water for a measurement in a Cochard-type rotor. We imagine that nano-particles occur at some moderate concentration in water (particles per ml) but not so common that they enter all cut open vessels in anyone measuring cycle, otherwise all vessels would embolize by the first or second spin.

## Results

### Vessel volumes and dimensions

Figure 2 shows the cross-sectional area of injected rubber in stem sections versus the distance from the injection surface to the cross-section of *Robinia* samples in Yangling. The dashed line represents the total volume of injected rubber computed from the integral of the best fit line times the distance. The arrows indicate the volume to the center of the small and large rotors. The total volume of all vessels can be estimated by the y-intercept of the solid-line plot times the stem length. For the stems lengths used in the two rotors, the values are 0.341 and 0.651 ml for the small and large stems, respectively. In comparison, the total water volume extracted into the cuvettes by 3 MPa tension was 0.83±0.07 (N=4) and 1.08±0.16 (N=16) ml for the small and large rotors, respectively. These extraction values are approximately double the volume of embolized vessels at 3 MPa; the first half of the extraction occurred from living cells before cavitation/loss of vessel water began (Du et al., 2019; Peng et al., 2019). During typical *K_h_* measurements, after a spin, the volume of water flowing into the vessels generally exceeded the likely volume of water-filled vessels by a factor of 2 to 4 except near the end of vulnerability curves when the volume of water needed to measure *K_h_* was nearly equal to the volume of water in the non-embolized vessels. So, at the end of each measuring cycle the water in the conducting vessel was replaced with ‘fresh water’; the significance of this will be address in the discussion.

**Fig. 2.**
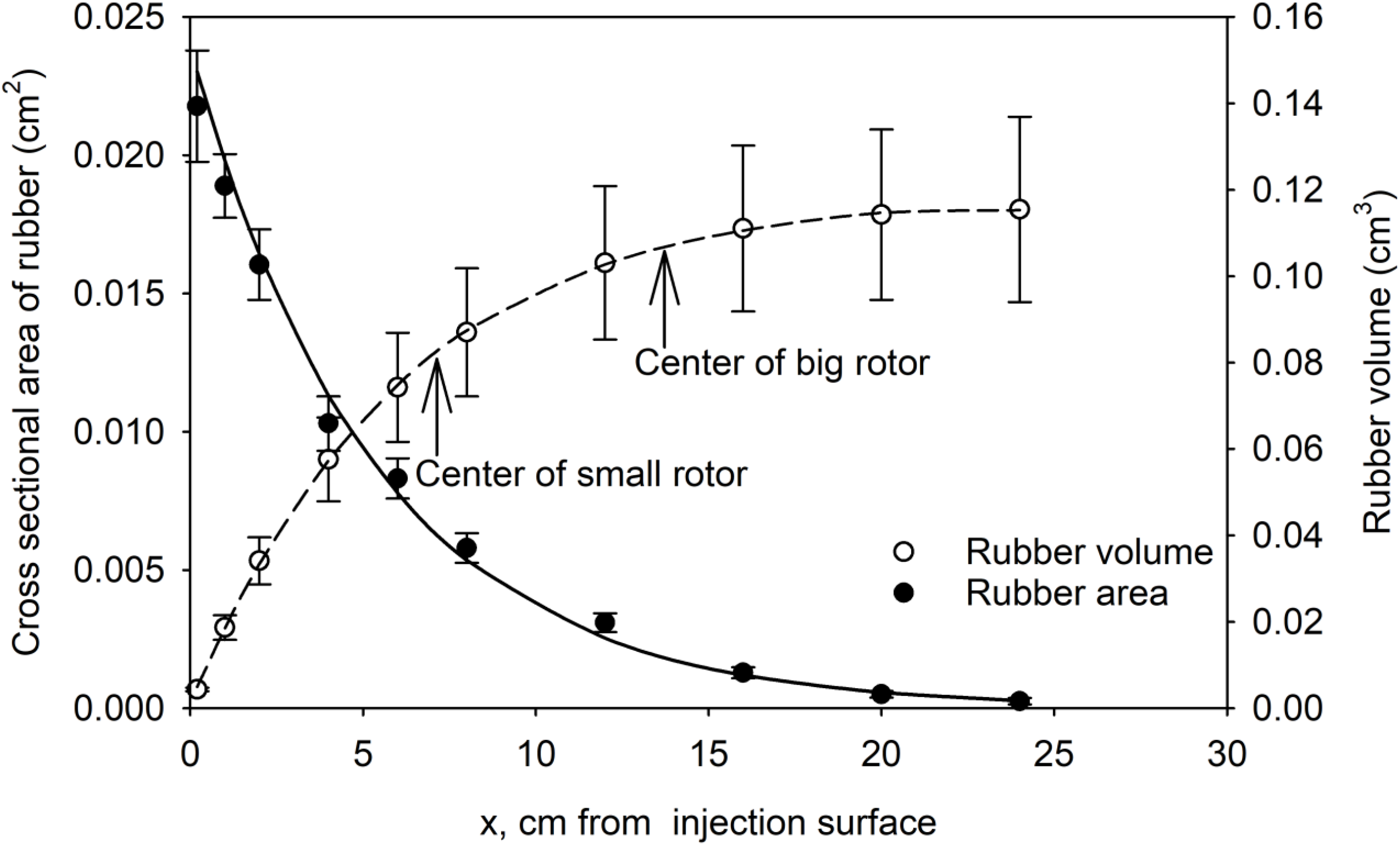
The left ordinate axis shows the cross-sectional area of rubber-filled vessels versus the distance from the injection surface (abscissa); points are the mean of measured values (*N* = 7 stems), and the solid line is the best fit curve: y = 2. 390E −02 exp(−0.187x), R^2^ = 0.9972. The right ordinate axis shows the volume of injected rubber from the injection surface (dashed line) given by the integral of the above equation. The volume is the maximum volume available to nano-particles that are filtered out by pit membranes between vessels.

Figure 3A demonstrates that the mean vessel diameter of the injected vessel increased with the distance from the injection surface of *Robinia* samples in Yangling. This is consistent with the notion that small-diameter vessels are shorter than larger-diameter vessels (Cai et al., 2010). The range of means was not large, i.e., from about 60 to 74 μm. Fig. 3B shows the mean vessel length versus the mean vessel diameter by bin size classes. Many of the short vessels would reach the axis of rotation of the small rotor, and the largest vessels would reach the axis of rotation of the large rotor. It is worth mentioning that vessel length in stem segments in *Robinia* change dramatically between trees or even between branches on the same tree (Wang et al. 2014), so we measured vessel length of the same stem segments that were spun in the Sperry rotors in this study.

**Fig. 3.**
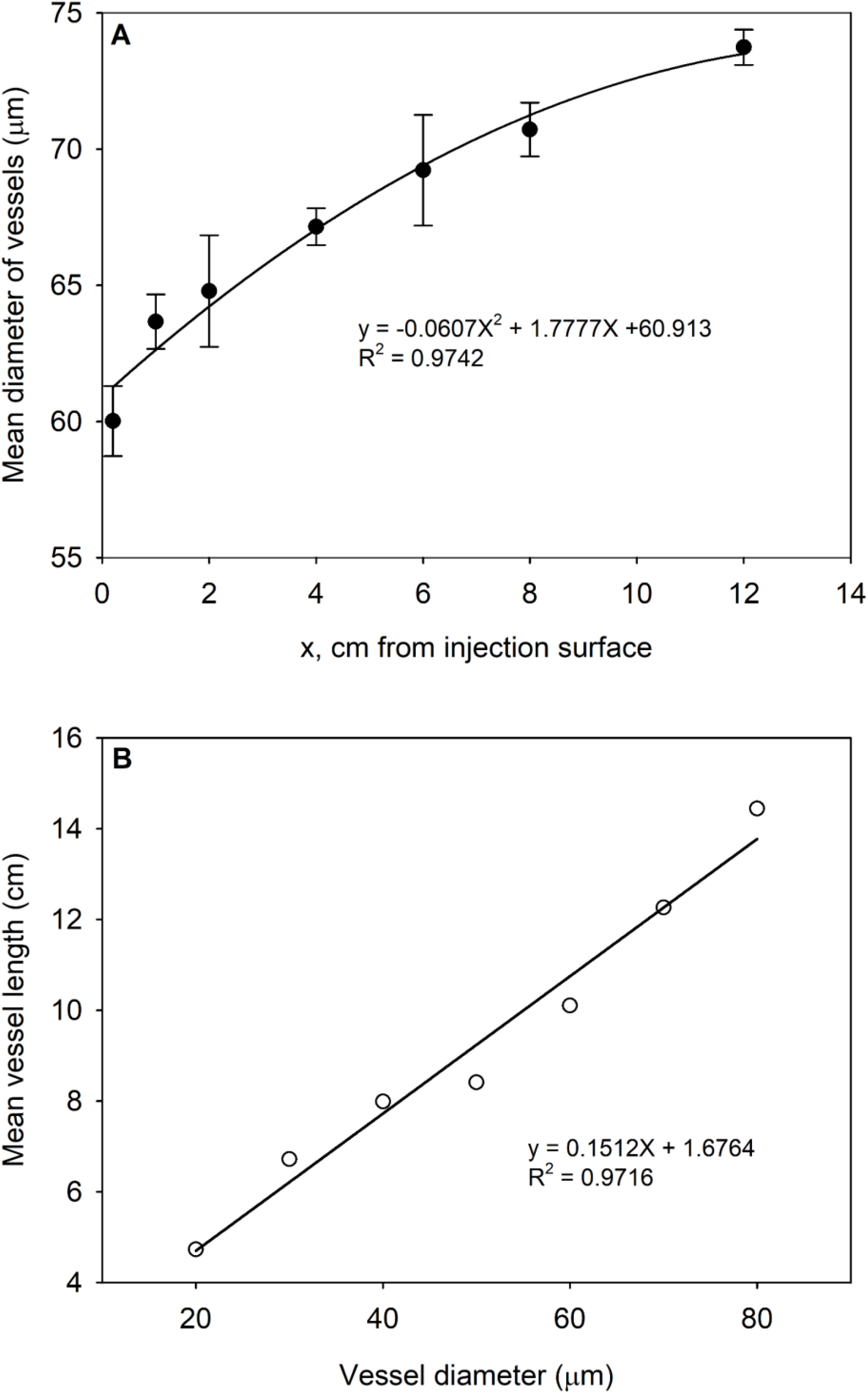
(A) Mean vessel diameter of rubber-injected vessels versus the distance from the injection surface. (B) Mean vessel length versus bin diameter size class. Note that mean vessel lengths are computed assuming that they randomly begin anywhere in the stem, so mean vessel lengths tend to exceed the distance of rubber infusion.

Mean vessel length and vessel diameter with standard error (SE) of all species were shown in Table 1. The results shown that *Acer* has the shortest vessel length (2.25 ± 0.09 cm) within all species, and it can be defined as short vessel species because it is shorter than radii of both big and small rotor (13.7 cm vs. 7.2 cm respectively). The mean vessel length of *Robinia* was close to the radius of big rotor, but larger than radius of small rotor. *Quercus* has the longest mean vessel length within all species, which was longer than the radius of big rotor. The vessel diameter comparison had the sequence as *Acer* < *Robinia* (Yangling) < *Robinia* (Jinhua), *Quercus*.

**Table 1.**
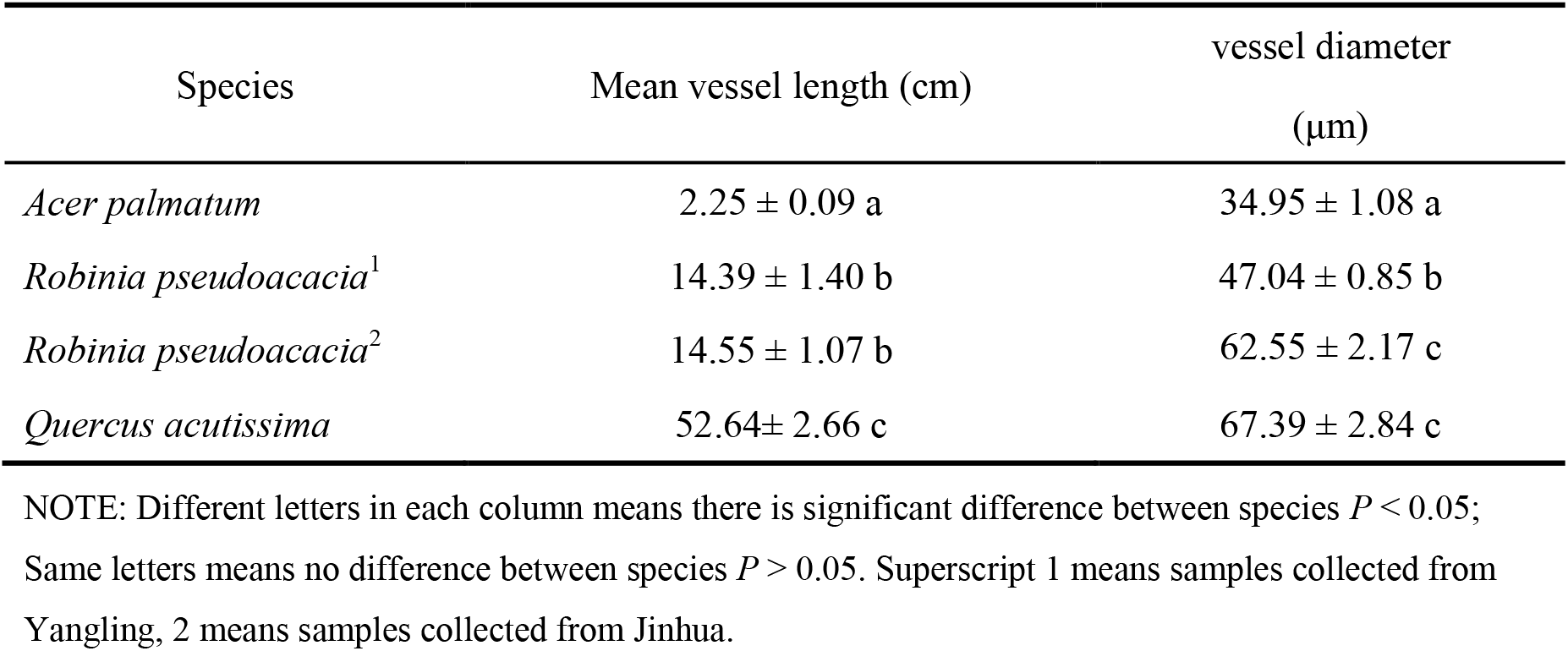
Mean ± SE of mean vessel length and vessel diameter of four species. Number of replicates: N = 6-9. Stems used for VCs measurement in large rotor.

### Early embolism in cut-open vessels spun repeatedly at the same tension

The purpose of this experiment was to test the underlying assumption of people who use rotors to spin stem segments at different tensions. In preliminary experiments when Sperry (Alder et al., 1997, Davis et al., 1999) tested their new rotor they determined how long the samples need to spin to get a stable result. But our results show that the answer turns out to be dependent on vessel length and when vessels are too long then quite unexpected results occur. In our *Robinia* experiments in Yangling, the same stem was repeatedly spun at 0.25 MPa tension, and after each spin, the *K_h_* was measured and the percentage loss of hydraulic conductivity, *PLC*, was computed. If the air-seeding hypothesis is the dominant mechanism of cavitation, then there should have been an initial drop in *PLC* after the first spin with little or no additional drop after successive spins. Contrary to this hypothesis, the *PLC* decreased by equal amounts between spins (Fig. 4A). This was further confirmed by the experiments in which the rotors were deliberately filled with unequal water masses (500 ± 1 mg for the small rotor and 2000 ± 1 mg for the large rotor). At the end of the spins, the water masses had equilibrated to within ±10 mg; hence, during the 6-min spin, an additional 0.5 or 2 g had flowed through the stem segments. The impact of the additional flow during the spin was to increase the decline of *K_h_/K_max_* after each spin. When the experiment in Fig. 4A was repeated in the large rotor (Fig. 4B), all the declined values of *K_h_/K_max_* were less than in the small rotor, both with and without equal masses of water in the cuvettes before the spin. The y-intercepts of both linear regressions in Fig. 4 were significantly less than 1 based on 95% confidence intervals (Table 3), which indicated that some additional nano-particles go into stem during flush rather than during spinning. The additional water passing through the sample *Robinia* segments (2 ml extra during each spin in large rotor) is similar to the amount of water passed through the segments when *K_h_* is measured in the LPFM, but the passage of water while spinning seems to increase the loss of *K_h_* after each spin (solid triangles versus open circles).

**Fig. 4.**
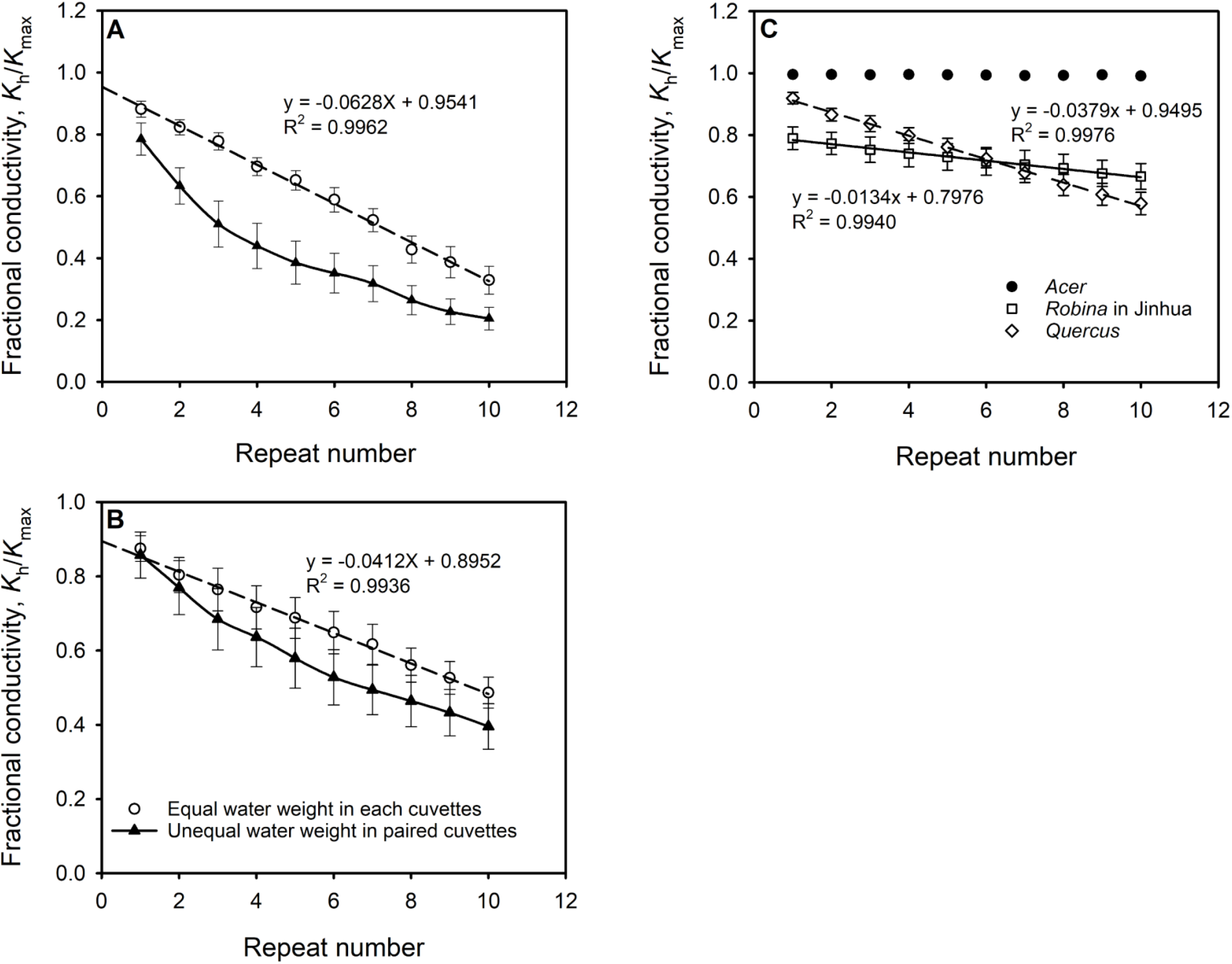
(A) *Robinia* stems of Yangling in the small rotor were subjected to 10 repeated spins of 6 min duration, and at the end of each spin, the hydraulic conductivity was measured. In the open circle points, the water mass in each cuvette was equal to ±1 mg or unequal by exactly 0.5 g (solid triangles). (B) The same as (A) except the stems were spun in the large rotor and the unequal masses of water were 2 g measured to ±1 mg. Stem lengths in the small and large rotors were 14.4 and 27.5 cm, respectively. (C) Comparisons of short to long vessel species measured in the large Sperry rotor in Jinhua.

We repeated these experiments years later at ZJNU with three species with short to long vessels relative to the diameter of the large Sperry rotor. These results are shown in Fig. 4C. *Robina* and *Quercus* both shown a significant decline of *K_h_/K_max_* with spin times, but in *Acer K_h_/K_max_* was constant. The vessel dimensions in these species were shown in Tables 1.

Location of living cells viewed in the cross section of *Acer, Robinia* and *Quercus* were shown in Fig. 5. Living cells (ray cells and xylem parenchyma) were stained black in the micrograph. All species have abundant ray cells near some of vessels, but *Quercus* and *Acer* have lots of xylem parenchyma compared to *Robinia*.

**Fig. 5.**
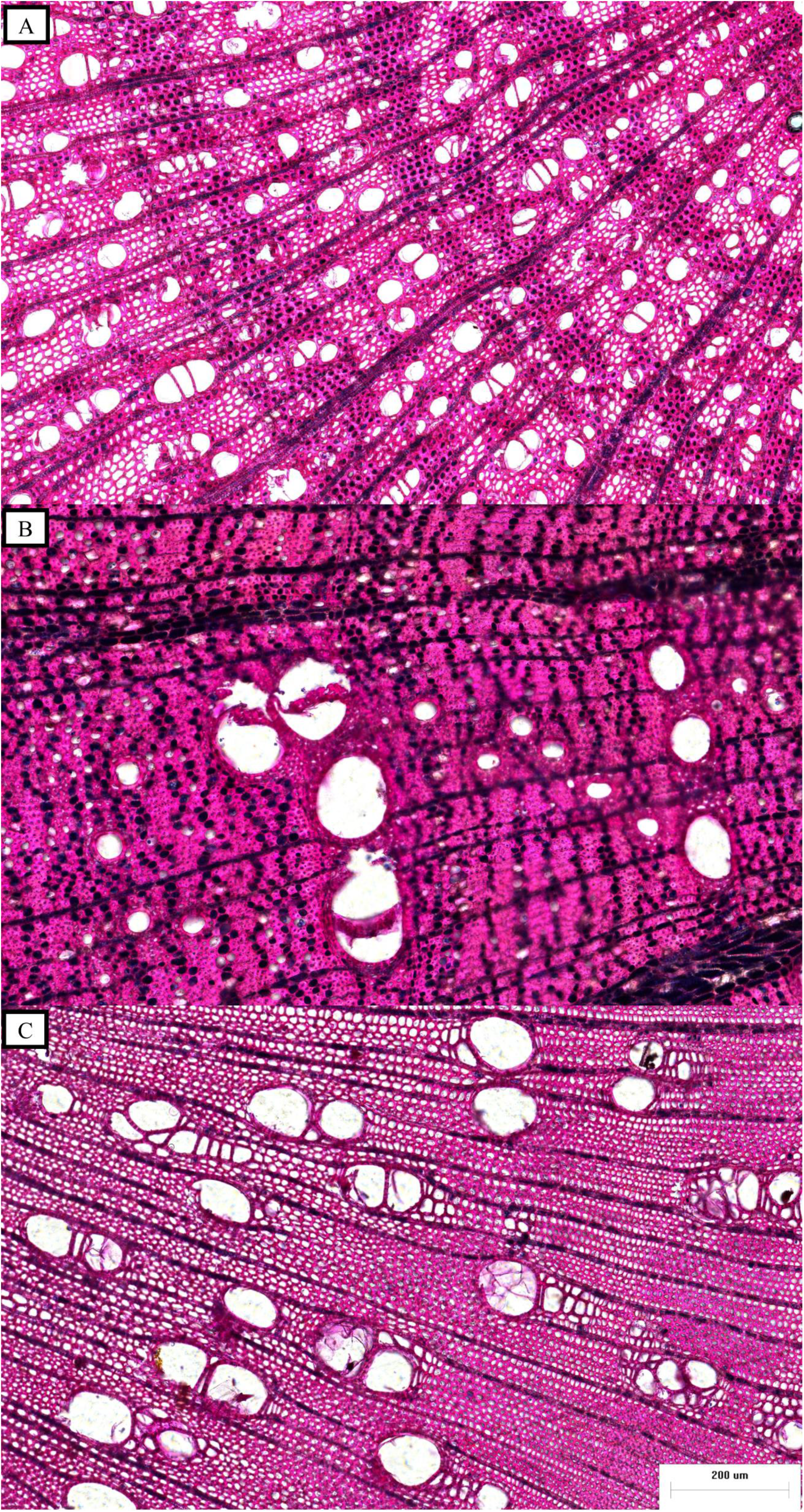
Starch in living cells stained with iodine in the cross section for (A) *Acer*, (B) *Quercus, and* (C) *Robinia*.

### T_50_ values measured by bench-top dehydration versus the Sperry rotor

Tentatively, Fig. 4 suggests that vulnerability curves, VC, of long-vessel species are determined by more than seeding of embolisms by the biggest pore out of all the pores in all the pit membranes of any given vessel. PLC is somehow the consequence of the volume of water that passes through the stem segments for long-vessel species but not for short-vessel species. This notion is also confirmed by the results in Fig. 6. Here we show that different vulnerability curves are obtained for *Robinia* segments depending on the method used. For the small rotor, if each segment is spun only once at any given tension (one-sample-one-tension method), *T_50_* = 2 MPa. In contrast *T_50_* = 1 MPa when each segment being spun 6 minutes and 10 different tensions (traditional method) in the small rotor (Fig. 6A). Similar differences were found in the large rotor (Fig. 6B).

**Fig. 6.**
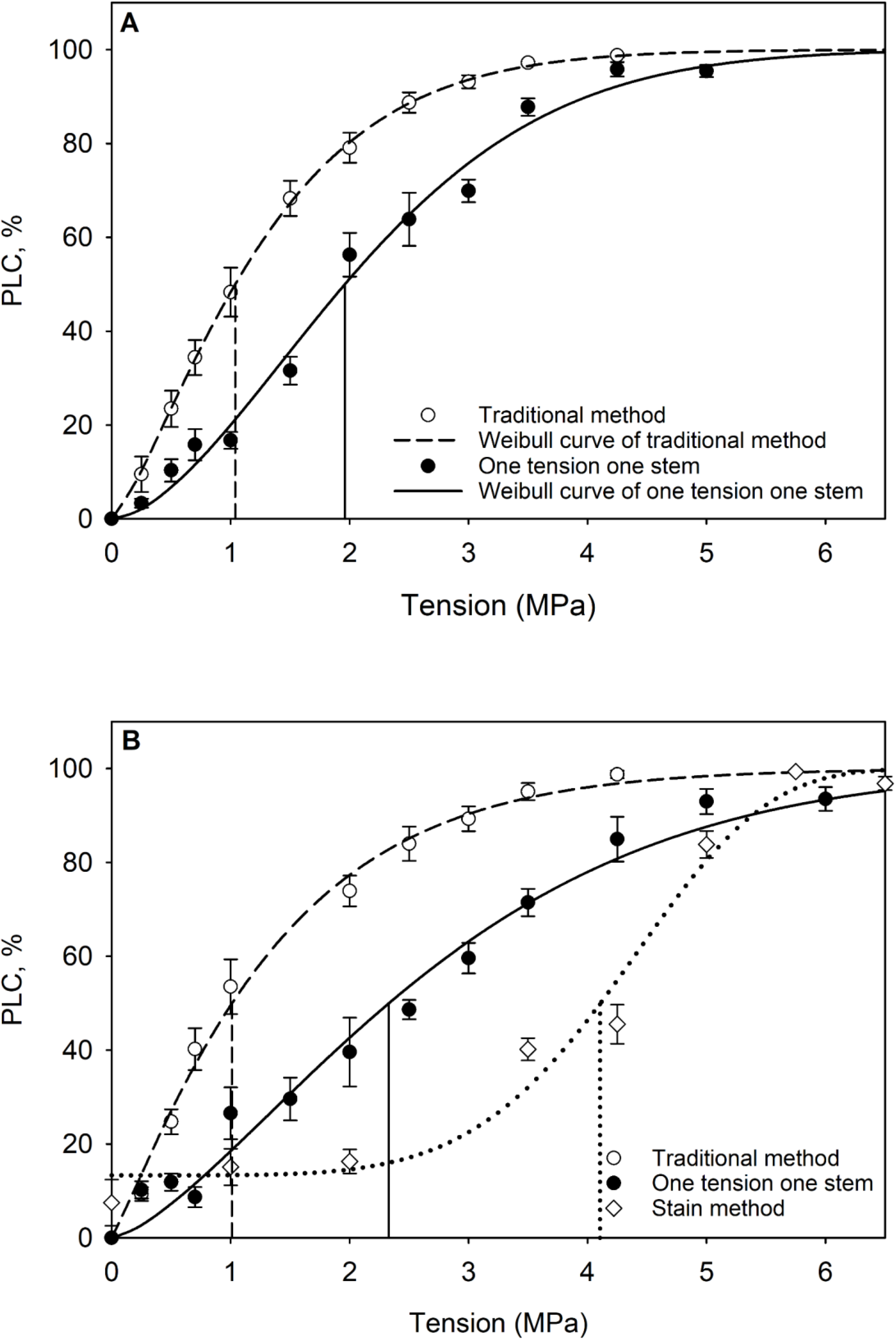
Vulnerability curves of *Robina* in Yangling by the traditional method and by the one-stem-one-tension method in (A) the small rotor, and (B) the large rotor. The vulnerability curves with the open circles and dashed line were obtained by the traditional way, where N=6 stems were centrifuged at each tension measured for PLC determination and then centrifuged again at each tension point on the curve. In the closed circles and solid line, different stems were spun at each tension and discarded after each measurement. The duration of centrifugation for each point was 6 minutes. The dotted line with diamonds in (B) gives the computed VC by the staining method evaluated at the axis of rotation without correction for ‘background’ embolism. Straight vertical lines shown *T*_50_ values for each method.

A plot of embolized vessels at the axis of rotation of big rotor visualized by staining (diamonds) is shown in Fig. 6B. Less than 10% of the vessel cross-sectional area was still cut open at the axis of rotation (see Fig. 2A solid line). Hence when the staining method is used to visualize PLC near the axis of rotation, less than 10% of the embolism from unstained versus stained vessels can be anomalous embolism due to cut open vessels. The *T_50_* values based on staining were significantly higher compared to other Sperry methods (*T_50_* = 4.1 ± 0.4 SE in staining method). This suggested that the higher PLC values in the circle points were caused by anomalous embolisms in cut-open vessels. Independently obtained staining experiments from our laboratory have confirmed that the % of embolized vessels falls off approximately linearly from the base to the apex (Yin et al., 2018). The % embolism observed from staining at tensions of 2, 3.5 and 4.25 MPa were all significantly lower than the open and closed circle points in Fig. 6B (all *P* < 0.01).

## Discussion

Many of the results above are similar to results obtained by Wang et al. (2014) using *Robinia* stem segments mounted in a Cochard rotor in a Chinese built ‘cavitron’, which has superior temperature control. Repeated measurements of *K_h_* were made on the same branch spinning at 900 RPM (*T* = 0.072 MPa), and Wang found that after many measurements over 200 minutes that *K_h_* reached a stable value (Circles in Fig. 7A), but in the first 25 minutes the *K_h_* fell linearly with time when measurements were done every 5 minutes (Squares in Fig. 7A). *Robinia* was a fortunate choice of species because there was a high diversity of vessel length from branch to branch. Hence within the same species Wang was able to study the impact of vessel length on the open vessel artifact. Fig. 7B, shows the impact of vessel length on stable *K_h_* after many measurements (similar to Fig. 7A) for about 200 minutes. These results for the Cochard rotor and the results in the Sperry rotor in this study suggest to us that the phenomenon is independent of the rotor type but strongly dependent on vessel length. One possible interpretation concerns hypothetical nano-particles.

**Fig. 7.**
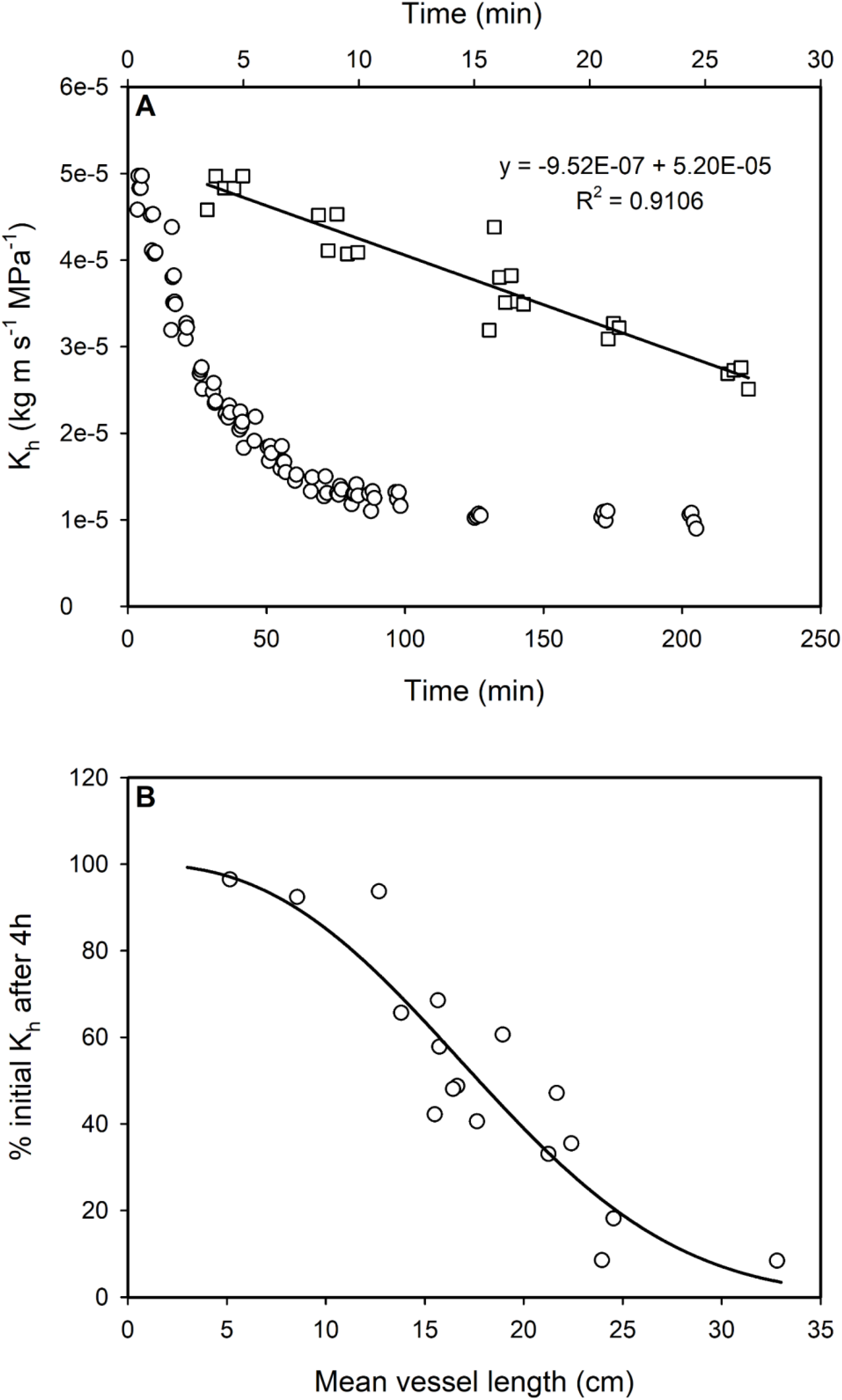
This is a replot and reanalysis of data from Wang *et al*. (2014) stem segment where hydraulic conductivity was repeatedly measured at low tension (0.072 MPa). (A) A typical example for a > 3 h of repeated measurements until a stable *K_h_* was observed (open circles); the squares are the same plot but for the first 30 minutes (the top x-axis). (B) The final *K_h_* is plotted as percentage of the initial *K_h_* on the y-axis versus the x-axis which plots the mean vessel length measured in the stem segment immediately after the *K_h_* values were measured. The stem segments lengths used were always 27.4 cm long; the trend line show a significant loss of *K_h_* whenever the mean vessel length was > about ¼ the stem segment length. All points are for *Robinia* which has a high variability of vessel length from between trees and even stems within a tree.

### The origin of hypothetical nano-particles

We had two hypotheses about the origin of the hypothetical nano-particles, i.e., laboratory water or cut stem surfaces. The first idea was that the nano-particles were present in laboratory water and would enter the vessels in proportion to the total volume flow. The flush introduced >100-fold more water volume into cut-open vessels than the typical volume flow during hydraulic conductivity measurements. If the origin of nano-particles was the filtered laboratory water, we would expect a disproportionate (> 100 times) more embolism in the first measurements in Fig. 4 than in subsequent measurements, but that was not found when repeated measurements of *PLC* were made at the same tension. The linear decrease between measurements was consistent with laboratory water being the origin of embolisms, but the y-intercept was inconsistent. The y-intercepts suggested a non-significant increase in early embolisms after the flush and the first spin compared to later spins, because the PLC changes between successive measurements were not significantly different from the uncertainty of the y-intercept.

Alternatively, living cells cut open near cut open vessels might be the source of nano-particles as these particles emerge at a constant rate. Figure 8 is a diagrammatic model (not to scale) explaining the origin of nano-particles from cut open living cells surrounded by semipermeable membranes (red lines). The osmotic pressure of the living cell contents is likely to be 1 to 3 MPa, but the osmotic pressure of the LPFM fluid is <0.1 MPa. Hence, osmotic water flow from the vessel across the membranes to the living cells (short yellow arrows) will push cell contents through the living cell lumen (long yellow arrows). The nano-particles (black circles) are presumed to always be present in the cytoplasm or are the degradation products of membrane-bound organelles that were pushed out by osmosis (yellow arrows) into the hypotonic water. Once organelles are in water, where water inflow will osmotically swell and burst the organelle. Water flow through the LPFM tubing (long black arrows) bends around low conductivity regions of the stem surface and then enters the vessel lumen as shown in Fig. 8 by the black arrows. This water flow near the surface picks up the cellular debris from the cut-open living cells and sweeps the debris into the vessel lumen. The discharge of nano-particles might last for hours. Membrane bound ‘organelles’ have been observed to emerge from cut stems and their swelling/shrinking and bursting due to changes in osmotic potential of bathing medium has been reported (Tyree et al., 1982); the primary purpose of Tyree et al. (1982) was to measure the kinetics of swelling and shrinking of emerging ‘organelles’, but MTT can confirm seeing such organelles breaking into components too small to see in a light microscope under oil-immersion optics when exposed to dilute solutions.

**Fig. 8.**
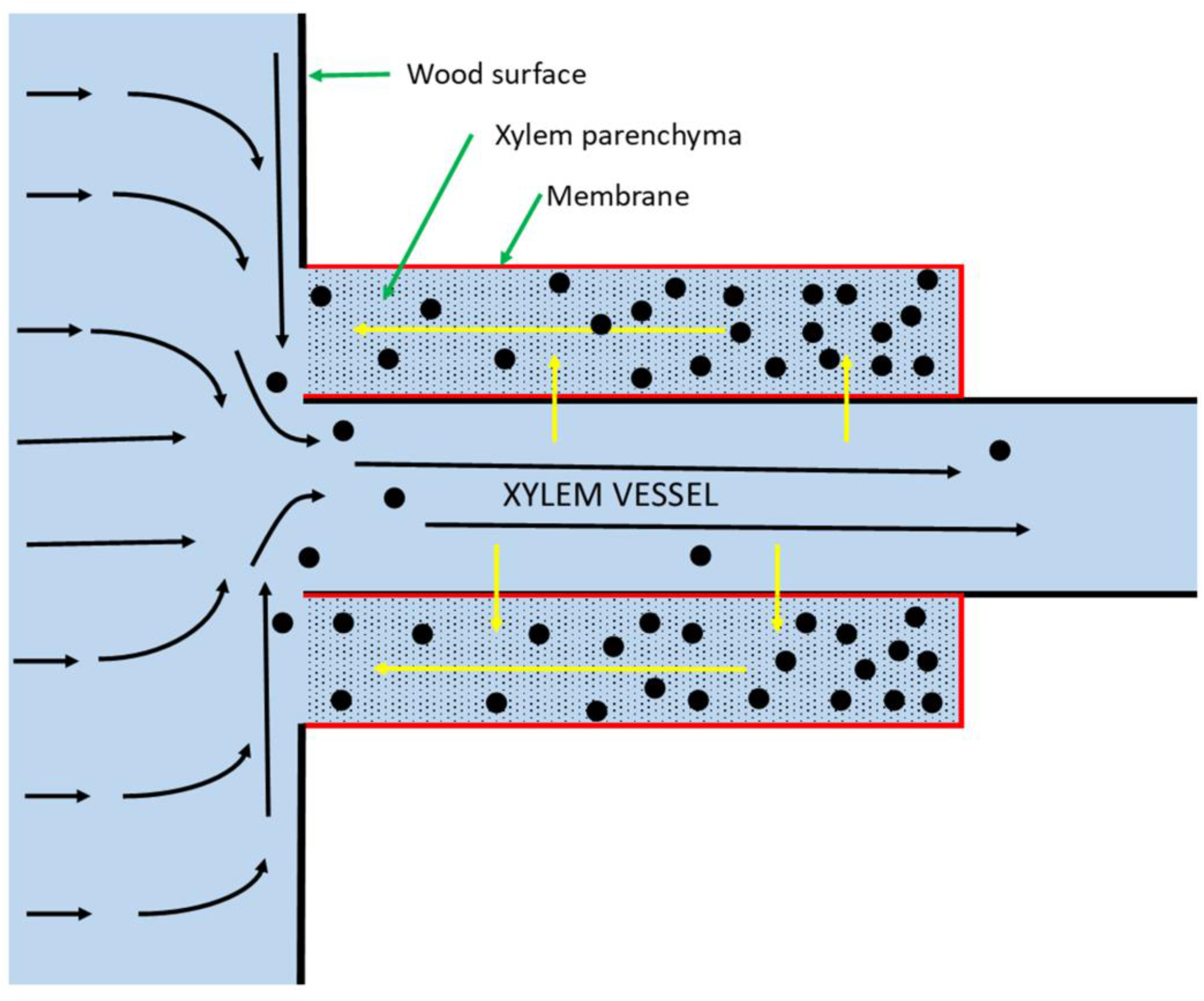
This is a radial view of a stem near the cross-sectional cut surface showing only one vessel and two living cells (wood fiber cells, etc., not shown). This is a diagrammatic representation of the origin of nano-particles (black circles) emerging from living cells cut open at the stem surface. Yellow arrows indicate osmotically induced flow, and black arrows indicate pressure-driven flow. See text for details.

The ideas presented about nano-particles are also consistent with the data in Fig. 4C. Our hypothesis is that the long-vessel artifact requires both long vessels and a source of nano-particles. *Robinia* and *Quercus* have both long vessels and living cells near to vessels and demonstrate the long-vessel artifact; but although *Acer* has lots of living cells nearby the vessels by vessels are short (Fig. 5). Hence, there is no early embolism in *Acer* (see Fig. 4C and Table 2). More experiments on up to 10 species with long- and short-vessel length are probably necessary to confirm our hypothesis; hence we hope our results will encourage studies of more species, beyond those we will do in the next year or two.

**Table 2.** The comparison of *T_50_* between bench-top dehydration and Sperry-type centrifugation methods in four species.

| Species | Bench-top dehydration | Centrifugation |
| --- | --- | --- |
| | 95% Confidence intervals | Mean ± SE |
| *Acer palmatum* | -3.41 (-3.66, -3.16) | - 3.24 ± 0.09 |
| *Robinia pseudoacacia*<sup>1</sup> | -3.74 (-3.66, -3.16) | - 1.01 ± 0.09 |
| *Robinia pseudoacacia*<sup>2</sup> | -2.25 (-2.47, -2.03) | - 1.39 ± 0.09 |
| *Quercus acutissima* | -2.11 (-2.29, -1.93) | - 1.03 ± 0.06 |

NOTE: Superscript 1 means samples collected from Yangling, 2 means samples collected from Jinhua.

**Table 3.**
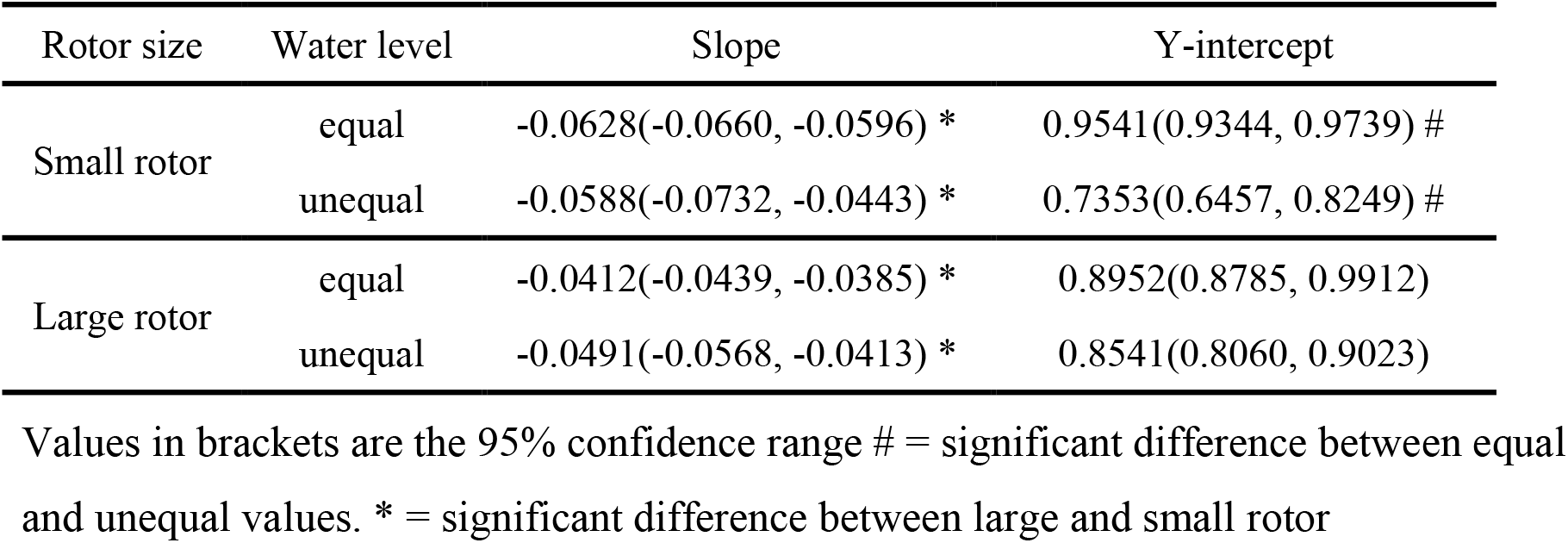
Statistical analysis for equal and unequal water level, repeat 10 cycles with the same tension.

| Rotor size | Water level | Slope | Y-intercept |
| --- | --- | --- | --- |
| Small rotor | equal | -0.0628(-0.0660, -0.0596) * | 0.9541(0.9344, 0.9739) # |
|  | unequal | -0.0588(-0.0732, -0.0443) * | 0.7353(0.6457, 0.8249) # |
| Large rotor | equal | -0.0412(-0.0439, -0.0385) * | 0.8952(0.8785, 0.9912) |
|  | unequal | -0.0491(-0.0568, -0.0413) * | 0.8541(0.8060, 0.9023) |

Values in brackets are the 95% confidence range # = significant difference between equal and unequal values. \* = significant difference between large and small rotor

### Does the rotor design make a difference?

The two centrifuge techniques in common use for measuring vulnerability curves differ primarily in their rotor designs, which determines when and where the *K_h_* values are measured. In the Sperry rotor, three stems can be spun simultaneously, but the stems must be removed from the rotor for measuring *K_h_* in a conductivity apparatus. Typically, the same stems are returned to spin at a higher tension. In the Cochard rotor, only one stem can be spun at a time, but *K_h_* can be measured while spinning. Typically, 0.3 to 0.7 g of water is injected into one cuvette depending on the optical system used to measure the movement of water from the injected cuvette. About two-thirds of the water injected into the cuvette moves through the stem while the *K_h_* measurements are made in the Cochard rotor and the rest passes through the stem while the stem spins at the next higher tension. In this study, the amount of water passing through the stem during *K_h_* measurements was a little less (0.15 to 0.4 g) but still sufficient to displace all the water in the cut-open vessels of *Robinia* (≤0.12 g, Fig. 2). Hence, neither rotor type avoids the hypothetical nano-particle problem.

Our hypothesis was that the water used to measure *K_h_* swept nano-particles into vessels that could induce embolism at quite low tensions. Hence, if repeated *K_h_* measurements were made at the same tension, then there should be a linear or curvilinear rate of decline in *K_h_* that depends on the number of measurements of *K_h_*, which injected new nano-particles. In the *Robinia* study of Yangling, the linear rate of decline in terms of *K_h_/K_max_* was 6% or 4% per spin for the small and large rotors, respectively, in equal water level experiment (circles in Fig. 4A, B). When the Sperry cuvettes were loaded with extra water in one end so that flow occurred during spinning, the rate of decline of *K_h_/K_max_* was increased initially to 15% and 9% per spin for the small and large rotors, respectively (triangles in Fig. 4A, B).

Our hypothesis was that nano-particles could potentially seed an embolism at a pressure below the air-seeding value in the pit membrane. If this is true, then there should be more embolisms on the upstream side of stem segments because nano-particles cannot pass through pit membranes. This has been confirmed by Yin et al. (2018); the number of embolized vessels was found in *Robinia* to linearly decrease with distance from the base to the apex after spinning stems to a tension of 1 MPa, which is sufficient to cause 50% PLC (Fig. 6). These findings reinforced the notion that nano-particles were filtered out by the pit membranes as water passed between vessels.

At the time of the experiments by Wang et al. (2014), our thinking was that the ‘nano-particles’ were in fact tiny air bubbles formed during injection of water into the source cuvette in the Cochard rotor. So, we redesigned the rotor to isolate the measuring cuvette from the injection cuvette by various techniques, but none of the techniques eliminated the anomalous early loss of *K_h_* at low tensions (Du et al., 2019). If the nano-particles are always present in laboratory water (even well-filtered water) or if the nano-particles are expelled from the cut-open living cells of recently cut stems, then the way to prevent the entry of nano-particles is to eliminate the *K_h_* measurements and measure instead the water extraction volume from embolized conduits. This approach gives more valid vulnerability curves (Pivovaroff et al., 2016; Peng et al., 2019), but the interpretation of the results in terms of PLC is complicated by the fact that part of the volume extraction is coming from tissues other than the lumens of xylem conduits.

It is now clear that vulnerability curves generated by both the Cochard rotor, and the Sperry rotor are subject to so-called ‘open vessel’ artifacts in long-vessel species (Cochard et al., 2013; Martin St-Paul et al., 2014; Torres-Ruiz et al., 2014; Wang et al., 2014). The nature of this artifact was consistent with the hypothesis that when vulnerability curves were measured on excised stems, a significant fraction of the embolisms was induced by a 5^th^ mechanism not previously considered by Tyree (1997) and Tyree and Zimmerman (2002), namely, early cavitation induced by nano-particles either generated at the cut surface of stems or always present in laboratory water. The data in this paper and in Wang et al. (2014) were entirely consistent with the hypotheses at the end of the introduction.

The so-called ‘gold standard’ of techniques for measuring vulnerability curves is the bench top dehydration method. This method works because whole shoots are harvested at a length equal to >2 times the maximum vessel length, and then cavitation is induced by slow bench top dehydration. The measurements of *K_h_* are made after dehydration to a tension typically measured with either a pressure chamber or a stem hygrometer. After the *K_h_* is measured on stem segments (far removed from the cut-open vessels at the base of the shoot), the stems are flushed to get maximum conductivity, *K_max_*, from which PLC is calculated from 100(1-*K_h_/K_m_*). The principal disadvantages of this method are: (1) that in some cases, the native PLC of some trees might be 10 to 40%, so bench top dehydration curves do not include the part of the vulnerability curve that is eliminated by native drought events and (2) the SEM values are often larger than in the centrifuge techniques thus making vulnerability curves quite inaccurate. The staining data in Fig. 6B were also a type of gold standard because these data were obtained totally by extracting water and then staining. Figure 9 is a replot of Fig. 6 but includes two recent vulnerability curves obtained by the benchtop dehydration method from our former laboratory in Northwest A&F University (Wang et al., 2014; Yin et al., 2018) as well as by the water extraction method (Peng et al., 2019).

**Fig. 9.**
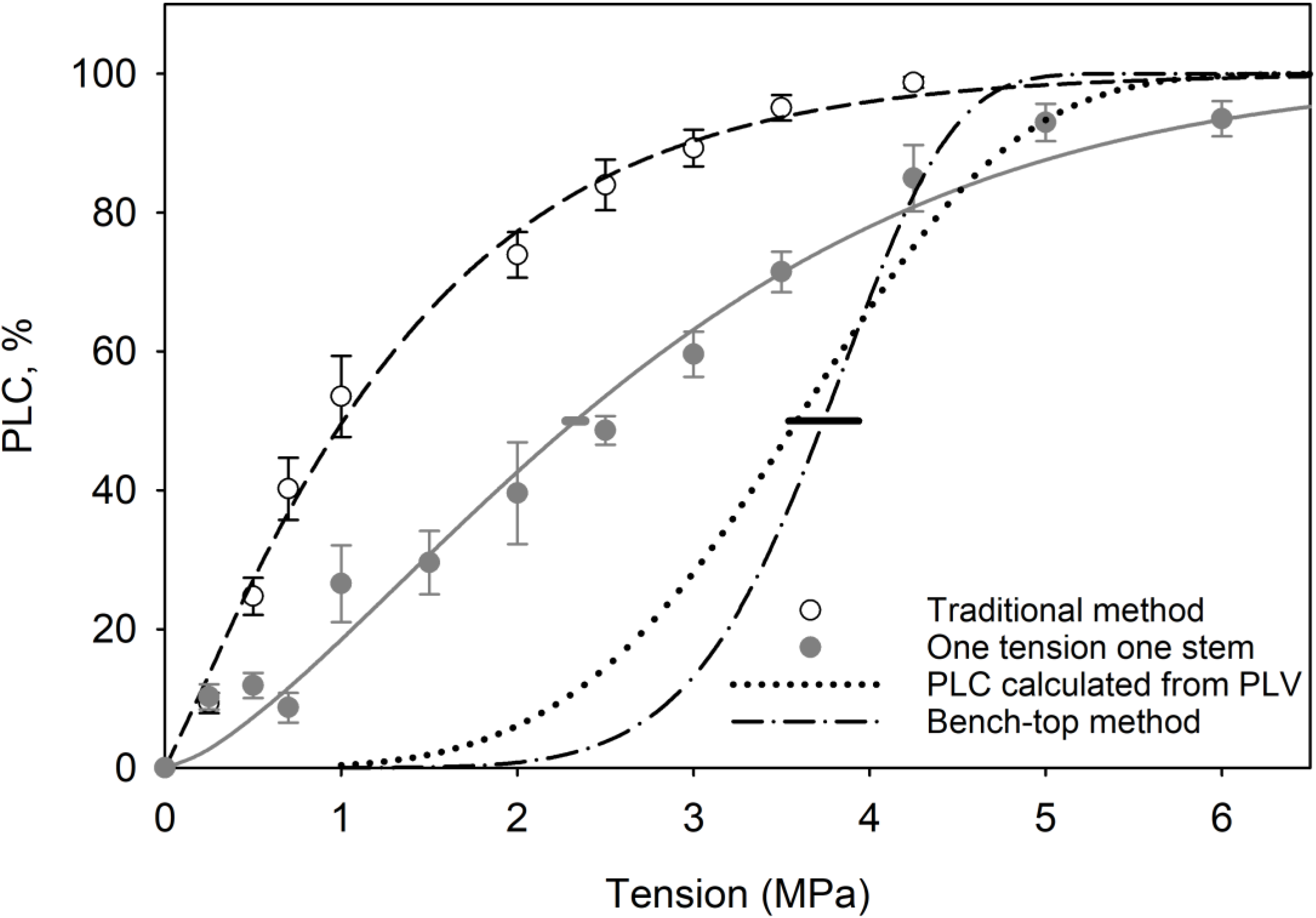
A replot of data collected using the large Sperry rotor (Fig. 4B) together with vulnerability curves collected by methods that should be immune to the early embolism by nano-particles. The dash-dotted line was obtained by the bench top dehydration method; stems are not flushed before dehydration, so no nano-particles are pushed into vessels (reproduced from Wang *et al*. 2014, also Yin *et al*. 2018 obtained similar results). The dotted line was obtained by the water extraction method (dotted Peng *et al*. 2019), which tends to push nano-particles away from the axis of rotation. Horizontal bars are 95% confidence interval of *T*_50_ by One tension one stem measurement in Sperry Centrifuge (gray) and Bench-top dehydration method (black) in Yangling.

The results of this study plus others from our laboratory (Wang et al., 2014; Du et al., 2019, Peng et al., 2019) suggest an urgent need to revisit the now-classic experiments by the Sperry group (Sperry et al., 2005; Wheeler et al., 2005; Hacke et al., 2006); these publications were the first to document well the relationships between xylem safety against cavitation (*T_50_* where a large value is a safe value) and various measurements of vessel diameter or cross-sectional area or parameters that arguably could be related to vessel size (such as xylem resistivity). The arguments about why the various parameters might correlate in the above papers are beyond the scope of this paper, but of the 27 species in the meta-analysis (Hacke et al., 2006), seven had vessel lengths >10 cm and two were slightly smaller but still close to the half-length of the stem segments in the small Sperry rotor used to determine *T_50_*-values. Of those long-vessel species, 7 species had *T_50_* values <2.8 MPa. In general, large diameter vessels tend to also be long vessels (Hacke et al., 2006; Cai and Tyree, 2014); hence, many *T_50_* values of large diameter vessel may be inaccurate and need to be re-measured to see if the conclusions of the three papers quoted from Sperry’s laboratory are still correct. What may be discovered is that the *T_50_* of species with big vessels has been underestimated, but new results may not improve any correlations between *T_50_* and measures of xylem hydraulic efficiency. A recent and much larger meta-analysis (Gleason et al., 2016) shows quite weak evidence of such tradeoffs. But the lack of correlations in Gleason et al. (2016) may change if a more reliable method, like the water extraction method (Peng et al., 2019) proves to work for a wide range of species.

## Conclusions

We wish to end with a positive note. Both the Sperry and Cochard rotors are valid techniques for measuring the VCs of species with short conduits. The criterion used for selection of short-vessel species might be that < 20% of the conductivity is due to vessels longer than the distance from the axis of rotation to the water levels in the cuvettes. This criterion is met by all conifers because of their tracheid lengths of less than 3 mm and by 65 to 75% of the short-vesselled angiosperm species studied so far. But the VCs of all species with long vessels need to be remeasured by advanced methods to see if the findings of this paper are generally true.

The two r-shaped curves on the left side of Fig. 9, compared to the s-shaped ‘normal’ curves, are often thought to be wrong and referred to as recalcitrant vulnerability curves caused by ‘open vessel artifacts.’ We now have a stronger reason to believe that the recalcitrant behavior is caused by nano-particles generated from the cut surface of stems (Fig. 8). But even if the open vessel artifact is NOT caused by nano-particles the results still stand. More than just tension, *T*, determines the PLC in vulnerability curves measured in centrifuges and something different happens during bench top dehydration. The way forward might be to remeasure VCs of all species with vessels longer than the radius of the rotors by using more reliable methods, for example the water extraction method (Peng et al. 2018). This may require years of work but might lead to results of more ecological significance than found in the literature meta-analysis by Gleason et al. (2016).

## Materials and methods

### Plant materials and sampling

In the summer of 2017, *Robinia pseudoacacia* L. current-year stem segments near the Weihe River in Yangling, Shaanxi, China (34°16′ N, 108°4′ E), were harvested from sun-exposed branches. Shoots 1- to 1.5-m long and with a 9-mm basal diameter were excised, sprayed with water, and then enclosed in black plastic bags with wet paper and transported to the laboratory within 0.5 h. The shoot base was recut under water; leaves and thorns were excised, and then the branches were rehydrated under water for 0.5 h. In 2019, *Acer palmatum* Thunb, *Robinia pseudoacacia* L., and *Quercus acutissima* Carruth, all collected at Zhejiang Normal University, Jinhua, China (29°07′ N, 119°38′ E) and used for the measurements done on *Robinia* collected in Yangling. The mean vessel length, vessel diameter, VCs with the large and small Sperry rotor, and VCs by benchtop dehydration were measured.

### Hydraulic conductivity measurements

Stem segments 275-mm or 144-mm long and 6.5 to 7.5 mm in diameter were recut under water, and then the scars from excised leaves and thorns were sealed by super glue to reduce evaporation. The stem segment ends were re-trimmed with a fresh razor blade and flushed with 0.01 M KCl (prepared with ultrapure water) at an applied pressure of 150-170 kPa pressure for 4-10 min for different species to ensure maximum hydraulic conductivity, *K_max_*.

After spinning stems in a centrifuge, the *K_h_* of stem segments was measured with a low-pressure flow meter (LPFM). We followed the protocol in Sperry et al. (2012) in which the background flow was measured immediately after the spin and again after the *K_h_* measurement. The background flow is attributed to tissue desiccation in the Sperry rotor, so the segments typically rehydrate while *K_h_* is measured. *K_h_* was always corrected for background flow, but usually the correction was less than 5%. The typical applied pressure was about 1.5 kPa (~15 cm of water head) for all species, which was low enough to avoid displacement of embolisms by moving water in vessels.

### Centrifuge measurements and vulnerability curves

Sperry rotors were used for inducing cavitation of stem segments; large and small rotors for 27.4 and 14.4 cm stems, respectively, were fabricated according to specifications in Alder et al. (1997). For *Robinia* experiment in Yangling, before and after spinning, the cuvettes were weighed to compute water extraction mass by difference. Before spinning, water was added gravimetrically to ± 1 mg with a syringe. This experimental design included equal and unequal water masses between the cuvettes. For the large rotor, 5 g of water was added in all cuvettes for the equal water level experiment, but 6 g versus 4 g was used for the unequal water level experiment in paired cuvettes. In the small rotor, 2 g of water was added in all cuvettes for equal water level experiments, but 2 g versus 1.5 g was used for unequal water level experiments in paired cuvettes. In 6 replicate experiments, equal water masses were used prior to each spin, and in 6 other replicates, unequal water masses were used as explained above during *Robinia* experiment in Yangling. The centrifuge was maintained at 25 °C, and all *K_h_* measurements were done at 25 °C.

In Jinhua, foam pads saturated with water were always used unless specifically stated otherwise in the results section, and the water mass was not measured. The pads were mounted in the vertical part of the cuvettes in the Sperry rotor. The function of the pads is to keep the cut surfaces of stem segments in better contact with water when the rotor was not spinning. While spinning, the pads are fully immersed in the water that has moved from the horizontal part of the cuvette to the vertical part by the centrifugal forces generated by the spin.

Both in Yangling and Jinhua, stems of all species were spun repeatedly at the same tension (0.25 MPa for 6 min) and removed for *K_h_* determination and then spun again at 0.25 MPa for a total of 10 cycles. In Jinhua, vulnerability curves were measured for 3 species with big rotors. In addition, repeated spins with increasing tension were used as in the standard Sperry protocol, and each tension was maintained for 6 minutes. In some cases, the same stem was used at each tension (standard Sperry protocol), and in other experiments, new stems were used at each tension, for a total of 60 stems to measure 10 different tensions (one-tension-one-stem).

In all experiments, the percentage loss of conductivity (*PLC*) = 100% (1 - *K*_h-correct_/*K*_max_), and a Weibull function was used to fit the relationship between the *PLC* and xylem tension: *PLC*/100 = 1 − exp [- (*T/B*)^*C*^], where *B* and *C* are curve fitting parameters, and they were obtained by minimizing the root mean square error (*RMS*_error_). The value of *T*_50_ was then calculated from *T*_50_ = *B [ln (2)]^1C^*.

### Vessel length and vessel diameter measurements

Mean vessel length was measured by the air-injection method (see details in Pan et al., 2014). In *Robinia* the same stems used for VCs were removed and used for air injection measurement. But for *Acer* and *Quercus* different stems with similar diameter were measured by air-injection methods. The vessel diameter of *Robinia* in Yangling were measured on cross sections by light microscopy using the same stems used for vessel length measurement.

#### Silicone rubber-injection method

Vessel length is frequently measured by a rubber injection method. The replacement of water with rubber is analogous to the replacement of water with air because pit membranes can stop both. Our primary purpose was to estimate the volume of a cut-open vessel, which equals the volume of injected rubber, but this method can also be used to obtain vessel length.

After injection and hardening of the silicone rubber, stems were cross-sectioned at several distances (from 0.2 to 27 cm) from the injection surface until less than 1 or 2% of the vessels were filled with rubber. Sections 18-μm thick were cut by a microtome (Leica RM 2235, Nussloch, Germany) and were mounted in water on a glass slide and placed on a microscope (Zeiss, Imager A.2 Göttingen, Germany) and photographed by UV light at 50× magnification for vessel cross-sectional area measurement and for the analysis described in detail in Cai et al., 2010; 2014; Pan et al., 2015; and Peng et al., 2019.

#### Vessel diameter

For *Robinia* in Yangling, the area of each rubber-filled vessel, *Ai*, was measured by Image analysis software (WinCELL 2013; Regent Instruments Inc) and then summed to get *Ar* at each distance *x*. For all species in Jinhua, the cross-section of each sample was stained with 0.02% (w/v) basic fuchsin dying solution (detailed see Peng et al., 2020), and finally each vessel area, *A_i_*, was measured by Image analysis software (WinCELL 2013; Regent Instruments Inc). Diameters were calculated from *D_i_*=*(4A_i_/π)^0.5^*.

### Visualization of embolized vessels by the stain method and corresponding VCs

In Yangling, *Robinia* stem segments used for staining were prepared using the same method as that for measuring vulnerability curves. Stems were spun for 1 h at various tensions to cavitate vessels. After spinning, the central 2-cm segment near the axis of rotation of the stem was cut under water by cutting progressively from each end to release tension until the last 2-cm segment was cut with a fresh razor blade. The staining method was like that described in detail by Wang et al. (2014), and Peng et al. (2019). After staining, segments were flushed with 0.01 M KCl at 130 kPa briefly to remove excess stain.

Then 18-μm cross-sections were sectioned from the middle of the 2-cm segments. Sections were mounted in glycerin on glass slides and photographed under a microscope (Zeiss, Imager A.2 Göttingen, Germany) at 50× magnification with a digital camera (Infinity 1-5C, Regent Instruments Inc., Quebec, Canada). Then, every stained (conducting) and unstained (embolized) vessel area (*A_v_*) in the whole cross-sections was measured by digital analysis software (Image-Pro Plus version 6.0). The few vessels with tyloses that were not included in the stained and unstained vessel areas.

Staining vulnerability curves were plotted based on the unstained vessel area and stained vessel area within the same cross-section at each tension. Percentage loss conductivity was computed from stained areas squared to give hydraulic weights, *PLC_s_*:

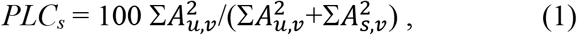

where *A_u,v_* = the area of the i^th^ unstained vessel, and *A_s,v_* = the area of the i^th^ stained vessel. Mean results were calculated from six to eight repeated stems.

### Presence of living cells near to vessels

Living cells were confirmed by iodine staining for starch, and lignified cell walls were stained by basic fuchsin in the cross section of stems. Cross sections thickness was approximately twice the presumed biggest living cells dimensions, which were determined based on tangential and cross sections. All cross sections were 25 μm thick for *Acer*, *Robinia* and *Quercus* and were mounted on slides, stained with iodine and basic fuchsin in sequence, viewed and photographed under a microscope (Leica, DM6B Wetzlar, Germany).

### Statistical analysis and replications

Student’s *t*-test was used for significance tests of PLC values at the given tension between traditional centrifuge method and one tension one stem method, and tests of *T*_50_ values of *Robinia* between traditional centrifuge method and theory calculation based on extracted method in Yangling. The 95% confidence intervals of *T*_50_ values in bench-top dehydration was calculated to compare with traditional Sperry centrifuge methods (Table 2). ANOVA were used for mean vessel length and mean vessel diameter comparisons among different species.

## Acknowledgements

P.G. and L.Z. thank the College of Forestry, Northwest A&F University, for laboratory space used during part of this research program, thank Yingjie Xiong, Xiaolin Wang for their assistance with anatomy measurements, also thank Dr. Yujie Wang providing helpful suggestions on writing. This research was supported by grants of the National Natural Science Foundation of China (No. 31770647), the Zhejiang Provincial Natural Science Foundation of China (LY19C150007).

